# Role of ventral subiculum neuronal ensembles in incubation of oxycodone craving after electric barrier-induced voluntary abstinence

**DOI:** 10.1101/2021.03.24.436801

**Authors:** Ida Fredriksson, Aniruddha Shekara, Sarah V. Applebey, Angelica Minier-Toribio, Lindsay Altidor, Carlo Cifani, Bruce T. Hope, Jennifer M. Bossert, Yavin Shaham

## Abstract

We recently developed a rat model of incubation of oxycodone craving where opioid seeking progressively increases after voluntary suppression of drug self-administration by adverse consequences of drug seeking. Here, we studied the role of ventral subiculum (vSub) neuronal ensembles in this incubation, using the activity marker Fos, muscimol-baclofen (GABAergic agonists) inactivation, and Daun02 chemogenetic inactivation.

We trained Sprague-Dawley or *Fos-lacZ* transgenic male and female rats to self-administer oxycodone (0.1 mg/kg/infusion, 6-h/d) for 14 days. The rats were then exposed for 14 days to an electric barrier of increasing intensity (0.1 to 0.4 mA) near the drug-paired lever that caused voluntary abstinence or were exposed to 14 days of forced abstinence. We tested Sprague-Dawley rats for relapse to oxycodone seeking without shock and drug on abstinence day 15 and extracted their brains for Fos-immunohistochemistry, or tested them after vSub vehicle or muscimol-baclofen injections on abstinence days 1 and 15. We performed Daun02 inactivation of relapse-activated vSub Fos neurons in Fos-lacZ transgenic rats on abstinence day 15 and then tested them for relapse on abstinence day 18.

Relapse after electric barrier-induced abstinence increased Fos expression in vSub. Muscimol-baclofen inactivation or Daun02 selective inactivation of vSub Fos-expressing neuronal ensembles decreased “incubated” oxycodone seeking after voluntary abstinence. Muscimol-baclofen vSub inactivation had no effect on non-incubated opioid seeking on abstinence day 1 or incubation after forced abstinence.

Our results demonstrate a selective role of vSub neuronal ensembles in incubation of opioid craving after cessation of drug self-administration by adverse consequences of drug seeking.

**Significance statement:** High relapse rate is a cardinal feature of opioid addiction and a major impediment for successful treatment. In humans, abstinence is often self-imposed, and relapse typically involves a conflict situation between the desire to experience the drug’s rewarding effects and negative consequences of drug seeking. To mimic this human condition, we recently introduced a rat model of incubation of oxycodone craving after electric barrier-induced voluntary abstinence. Here, we used the activity marker Fos, muscimol-baclofen (GABAergic agonists) inactivation, and Daun02 chemogenetic inactivation to demonstrate a selective role of vSub neuronal ensembles in incubation of oxycodone craving after electric barrier-induced voluntary abstinence, but not in incubation of opioid craving after forced abstinence or non-incubated opioid seeking during early abstinence.

## Introduction

High relapse rates are a major contributor to the opioid crisis (Rudd et al., 2016; Nunes et al., 2018). In humans, relapse and craving are commonly triggered by reexposure to cues and contexts previously associated with drug use (Wikler, 1973; O’Brien et al., 1992; Sinha, 2011). In laboratory rats, opioid seeking progressively increases or incubates after homecage forced abstinence from heroin (Shalev et al., 2001; Theberge et al., 2012) and oxycodone (Blackwood et al., 2019; Fredriksson et al., 2020) self-administration. Over the last two decades, mechanistic studies have primarily focused on incubation of psychostimulant craving (Pickens et al., 2011; Wolf, 2016; Dong et al., 2017). In contrast, the brain mechanisms of incubation of opioid craving are largely unknown (Reiner et al., 2019).

In the classical incubation of craving rat model, drug seeking is assessed after homecage forced abstinence (Grimm et al., 2001; Venniro et al., 2016). This contrasts with the human condition, where abstinence is often self-imposed and relapse episodes typically involve conflict situations where the individual using drug chooses between experiencing the drug’s rewarding effect and the potential adverse consequences of pursuing the drug (Marlatt, 1996; Epstein and Preston, 2003). Based on these considerations, we recently developed a rat conflict model of incubation of craving after voluntary abstinence, achieved by introducing an “electric barrier” near the drug-paired lever that the rats must cross to gain access to the self-administered drug. Using this model, we reported that oxycodone seeking is higher in the relapse tests after 15 and 30 abstinence days than after 1 day, demonstrating incubation of oxycodone craving after electric barrier-induced abstinence (Fredriksson et al., 2020). Unexpectedly, in both sexes incubation of oxycodone craving was stronger after voluntary abstinence than after homecage forced abstinence (Fredriksson et al., 2020).

The goal of the present study was to begin characterizing the mechanisms of incubation of oxycodone craving after electric barrier-induced voluntary abstinence by determining the role of ventral subiculum (vSub) in this incubation. We focused on vSub because our previous studies demonstrate a role of vSub in relapse to alcohol seeking after punishment-induced abstinence (Marchant et al., 2016b) and context-induced reinstatement of heroin seeking (Bossert and Stern, 2014; Bossert et al., 2016). There is also evidence for a role of vSub in approach-avoidance conflict decision-making (Ito and Lee, 2016; Schumacher et al., 2018).

We first used immunohistochemistry for the activity marker Fos (Morgan and Curran, 1991) to assess whether incubation of oxycodone craving after electric barrier-induced abstinence is associated with increased activity in vSub. Next, we used the muscimol-baclofen (GABAA and GABAB receptor agonists) reversible inactivation procedure (McFarland and Kalivas, 2001) to determine the general role of vSub in incubation of oxycodone craving after either electric barrier-induced abstinence or forced abstinence. Finally, we used the Daun02 inactivation method (Koya et al., 2009) to determine the specific role of vSub *neuronal ensembles* in incubation of oxycodone craving after electric barrier-induced abstinence. This method was developed to study the causal role of Fos-expressing neuronal ensembles in learned behaviors (Koya et al., 2009; Cruz et al., 2013). In *Fos-lacZ* transgenic rats (Kasof et al., 1996), both Fos and beta-galactosidase (β-gal) are induced in neurons that are strongly activated during learned behaviors. The prodrug Daun02 is injected into discrete brain areas 90 min later. β-gal converts Daun02 into daunorubicin only in the activated neurons, which inactivates and kills these neurons (Pfarr et al., 2015; Engeln et al., 2016). We and others have previously used the Daun02 inactivation method to assess causal roles of neuronal ensembles in different brain areas in context-induced reinstatement of cocaine and heroin seeking (Bossert et al., 2011; Cruz et al., 2014), incubation of drug craving (Fanous et al., 2012; Funk et al., 2016; Caprioli et al., 2017), and cocaine and alcohol seeking (Pfarr et al., 2015; de Guglielmo et al., 2016; Laque et al., 2019; Warren et al., 2019; Kane et al., 2020).

## Materials and Methods

### Subjects

In Exp. 1-3, we used male (n=68) and female (n=55) Sprague‐Dawley rats (Charles River) weighing 270-390 g and 180-240 g, respectively, prior to surgery. In Exp. 4, we used male (n=40) and female (n=46) *Fos-lacZ* transgenic rats (Koya et al., 2009), weighing 300-530 g and 180-320 g, respectively, prior to surgery. We maintained the rats under a reverse 12:12 h light/dark cycle (8:00 a.m. lights off) with food and water freely available. We housed two rats/cage prior to surgery and then individually after surgery. We performed the experiments in accordance with NIH Guide for the Care and Use of Laboratory Animals (8th edition), under a protocol approved by the local ACUC. We excluded 55 of the 209 rats used in the study due to catheter failure (n=1), failure to acquire oxycodone self-administration (n=2), poor health (n=19), cannula misplacement (n=18), loose head caps/damaged brains (n=12), reaction to intracranial injection (n=1), or outliers during the relapse tests > 3 STD (n=2).

### Drugs

We received oxycodone hydrochloride (HCl) from NIDA pharmacy and dissolved it in sterile saline. We chose a unit dose of 0.1 mg/kg for self-administration training based on our previous studies (Bossert et al., 2019; Fredriksson et al., 2020). In Exp. 2-3, we dissolved muscimol-baclofen (Tocris Bioscience) in sterile saline and injected it intracranially at a dose of 50+50 ng in 0.5 μl/side (Stopper and Floresco, 2014; Venniro et al., 2017; Reiner et al., 2020) 15-30 min before the relapse test sessions. In Exp. 4, we dissolved Daun02 (Sequoia Research Products) in vehicle solution containing 5% DMSO, 6% Tween-80, and 89% 0.1 M sterile PBS and injected it intracranially at a dose of 4 μg in 1.0 μl/side 90 min after the induction session. We chose the Daun02 concentration based on our previous study (Caprioli et al., 2017).

### Intravenous surgery

We anesthetized the rats with isoflurane (5% induction; 2–3% maintenance, Covetrus). We attached silastic catheters to a modified 22‐gauge cannula cemented to polypropylene mesh (Amazon or Industrial Netting), inserted the catheter into the jugular vein and fixed the mesh to the mid‐scapular region of the rat (Caprioli et al., 2015; Caprioli et al., 2017; Venniro et al., 2018). We injected the rats with ketoprofen (2.5 mg/kg, s.c., Covetrus) after surgery and the following day to relieve pain and decrease inflammation. We allowed the rats to recover for 6–8 days before oxycodone self‐administration training. During recovery and all experimental phases, we flushed the catheters every 24-48 h with gentamicin (4.25 mg/ml, Fresenius Kabi, USA) dissolved in sterile saline. If we suspected catheter failure during training, we tested patency with the short-acting barbiturate anesthetic Brevital (methohexital sodium, Covetrus; 10 mg/ml in sterile saline, 0.1–0.2 ml injection volume, i.v.) or Diprivan (propofol, NIDA pharmacy, 10 mg/ml, 0.1-0.2 ml injection volume, i.v.), and if not patent, we re-catheterized the other jugular vein and continued training the next day or eliminated the rat from the study.

### Intracranial surgery

We performed the intracranial surgery at the same time of the intravenous surgery for all experiments except for the rats in Exp. 2 in which we tested for relapse on abstinence day 15. For these rats, we performed the intracranial surgery one day after the last day of oxycodone self-administration training. We gave the rats 3 days to recover before the voluntary abstinence (electric barrier) phase (see below). We anesthetized the rats and, using a stereotaxic instrument (Kopf), implanted bilateral guide cannulas (23 gauge; Plastics One) 1 mm above the vSub. We set the nose bar at −3.3 mm and used the following coordinates from bregma: anteroposterior (AP), −6.0 mm; mediolateral (ML), ±5.3 mm (4° angle); dorsoventral (DV), −7.5 mm for males and −7.2-7.5 mm for females. We anchored the cannulas to the skull with jeweler’s screws and dental cement. We used the above coordinates based on our previous studies (Bossert and Stern, 2014; Bossert et al., 2016; Marchant et al., 2016b).

### Intracranial injections

Four days before the intracranial injections, we habituated the rats to the injection procedure. Habituation consisted of three phases. We first exposed the rats to the injection cage (an empty cage containing bedding). The following day, we gently removed the cannula blockers before exposing the rats to the injection cage. On the last day of habituation, we gently lowered down the injectors and placed them in the injection cage. On the test day, we connected the syringe pump (Harvard Apparatus) to 10 μl Hamilton syringes and attached the Hamilton syringes to the 30-gauge injectors via polyethylene-50 tubing; the injectors were extended 1 mm below the tips of the guide cannulas. In Exp. 2-3, we injected vehicle (saline) or muscimol-baclofen (50+50 ng in 0.5 μl/side) at a rate of 0.5 μl/min and left the injector in place for an additional minute to allow diffusion. After testing, we deeply anesthetized the rats with isoflurane and removed their brains and stored them in 10% formalin. We sectioned brains at 50 μm using a Leica Microsystems cryostat and stained sections with cresyl violet to verify the placement of the cannulas. In Exp. 4, we injected vehicle or Daun02 (4 μg/1.0 μl/side) at a rate of 0.5 μl/min and left the injectors in place for 2 additional min to allow diffusion.

### Fos immunohistochemistry

We based our Fos immunohistochemistry procedure on our previous reports (Bossert et al., 2012; Bossert et al., 2016). Ninety minutes after the relapse test in Exp. 1 and Exp. 4, we deeply anesthetized the rats with isoflurane (~80 s) and perfused them transcardially with 1∼100 ml of 0.1 m PBS, pH 7.4, followed by ∼400 ml of 4% paraformaldehyde in PBS. In Exp. 1, we also perfused the No‐test rats (taken from their homecage) the day following the relapse test. We removed and post‐fixed the brains in 4% paraformaldehyde for 2 h before transferring them to 30% sucrose in PBS for 48 h at 4°C. We subsequently froze the brains in powdered dry ice and stored them at −80°C until sectioning. We cut coronal sections (40 μm) containing the vSub using a cryostat (Leica Microsystems). We divided the sections into five series (200 μm apart), collected them in PBS containing 0.1% sodium azide and stored them at 4°C.

We rinsed free‐floating sections in PBS (3×10 minutes), incubated them for 1 h in 4% bovine serum albumin (BSA) in PBS with 0.3% Triton X‐100 (PBS‐TX) and incubated them overnight at 4°C with rabbit anti‐c‐Fos primary antibody [Phospho‐c‐Fos (Ser32), Cell Signaling Tech, RRID: AB_2247211, D82C12 diluted 1:8000] in 4% BSA in 0.3% PBS‐TX. We then rinsed the sections in PBS and incubated them for 2 h with biotinylated anti‐rabbit IgG secondary antibody (BA‐1000, Vector Laboratories) diluted 1:600 in 4% BSA in 0.4% PBS‐TX. We rinsed the sections again in PBS and incubated them in avidin–biotin– peroxidase complex (ABC Elite kit, PK‐6100, Vector Laboratories) in 0.5% PBS‐TX for 1 h. We then rinsed the sections in PBS, developed them in 3,3′‐diaminobenzidine, rinsed them in PBS, mounted them onto chrome alum/gelatin‐coated slides and air dried them.

We dehydrated the slides through a graded series of alcohol concentrations (30, 60, 90, 95, 2 × 100% ethanol), cleared with Citra Solv (Fisher Scientific) and coverslipped them with Permount (Fisher Scientific). We captured bright-field images of vSub with a Retiga 2000R CCD camera (QImaging) attached to a Zeiss microscope Axio Scope A1 using a 10X objective. We counted Fos-immunoreactive (IR) nuclei, characterized by brown nuclear staining, using iVision (4.5.0, Biovision Technologies). For each rat, we quantified cells in both hemispheres of two sections and computed a mean of these counts per area. We captured and quantified the following Bregma coordinates: −5.5 to −6.3 mm for vSub. We performed image-based quantification of Fos-labeled neurons in a blind manner by two independent observers (inter-rater reliability for counting between IF and CC, r=0.99, p>0.01 and IF and AS r=0.99, p, >0.01, for Exp. 1 and 4, respectively).

### X-gal histochemistry for β-gal visualization in *Fos-lacZ* rats

The X-gal assay is based on our previous studies (Koya et al., 2009; Bossert et al., 2011; Warren et al., 2016). Ninety minutes after the relapse tests in Exp. 4, we anesthetized the rats with isoflurane and perfused them transcardially with ∼100 ml of 0.1 m PBS, pH 7.4, followed by ∼400 ml of 4% paraformaldehyde in PBS. We removed the brains and postfixed them in 4% paraformaldehyde for 2 h before transferring them to 30% sucrose in PBS for 48 h at 4°C. We froze the brains in dry ice and stored them at −80°C. We collected coronal brain sections (40 μm) of vSub in PBS containing 0.1% sodium azide and stored them at 4°C until further processing.

We washed free-floating sections three times for 10 min each in PBS and incubated them in reaction buffer (2.4 mM X-gal, 100 mM sodium phosphate, 100 mM sodium chloride, 5 mM EGTA, 2 mM MgCl2, 0.2% Triton X-100, 5 mM K3FeCN6, and 5 mM K4FeCN6) for 6.5 h at 37°C with gentle shaking. We washed sections three times for 10 min each in PBS and mounted them onto chrom-alum/gelatin-coated slides and air dried. We dehydrated the slides through a graded series of alcohol (30%, 60%, 90%, 95%, 2 × 100% ethanol), cleared them with Citrasolv, and coverslipped the slides with Permount. We captured bright-field images of vSub with a Retiga 2000R CCD camera (QImaging) attached to a Zeiss microscope Axio Scope A1 using a 10X objective. We counted β-gal-expressing nuclei, characterized by blue nuclear staining, using iVision (4.5.0, Biovision Technologies) in sampling areas around vSub injection site (left and right hemispheres) in 4 coronal sections per rat. We performed image-based quantification of the β-gal-labeled neurons in a blind manner by two independent observers (inter-rater reliability of IF and AS, r=0.99, p>0.01).

### Self-administration apparatus

We used Med Associates chambers. Each chamber had two levers located 7.5–8 cm above the grid floor on opposing walls. Responding on the active retractable lever activated the infusion pump, while lever presses on the inactive, non‐retractable, lever had no consequences. We equipped each chamber with a stainless-steel grid floor connected to a shocker (Med Associates ENV-410B).

### General behavior procedure

The experiments generally consisted of some or all of the following phases: oxycodone self‐administration training (14 days), early tests for oxycodone seeking (abstinence day 1), electric barrier-induced abstinence (13 or 16 days), and late tests for oxycodone seeking (abstinence day 15 or 18). We provide details of the different phases for each experiment below.

### Oxycodone self-administration training

We trained the rats to self‐administer oxycodone‐HCl for 6-h/day (six 1‐h sessions separated by 10 min) for 14 days. Oxycodone was infused at a volume of 100 µl over 3.5 s at a unit dose of 0.1 mg/kg/infusion. Each session began with illumination of a red houselight that remained on for the entire session, followed 10 s later by the insertion of the active lever. Active lever presses led to oxycodone infusions that were paired with a 20-s tone‐light cue under a fixed-ratio 1 (FR1) reinforcement schedule (fixed interval 20-s [FI20] schedule). At the end of each 1-h session, the houselight turned off and the active lever retracted. We limited oxycodone intake to 15 infusions per h. We show the training data of Exp. 1-4 in Fig. 1.

**Figure 1.**
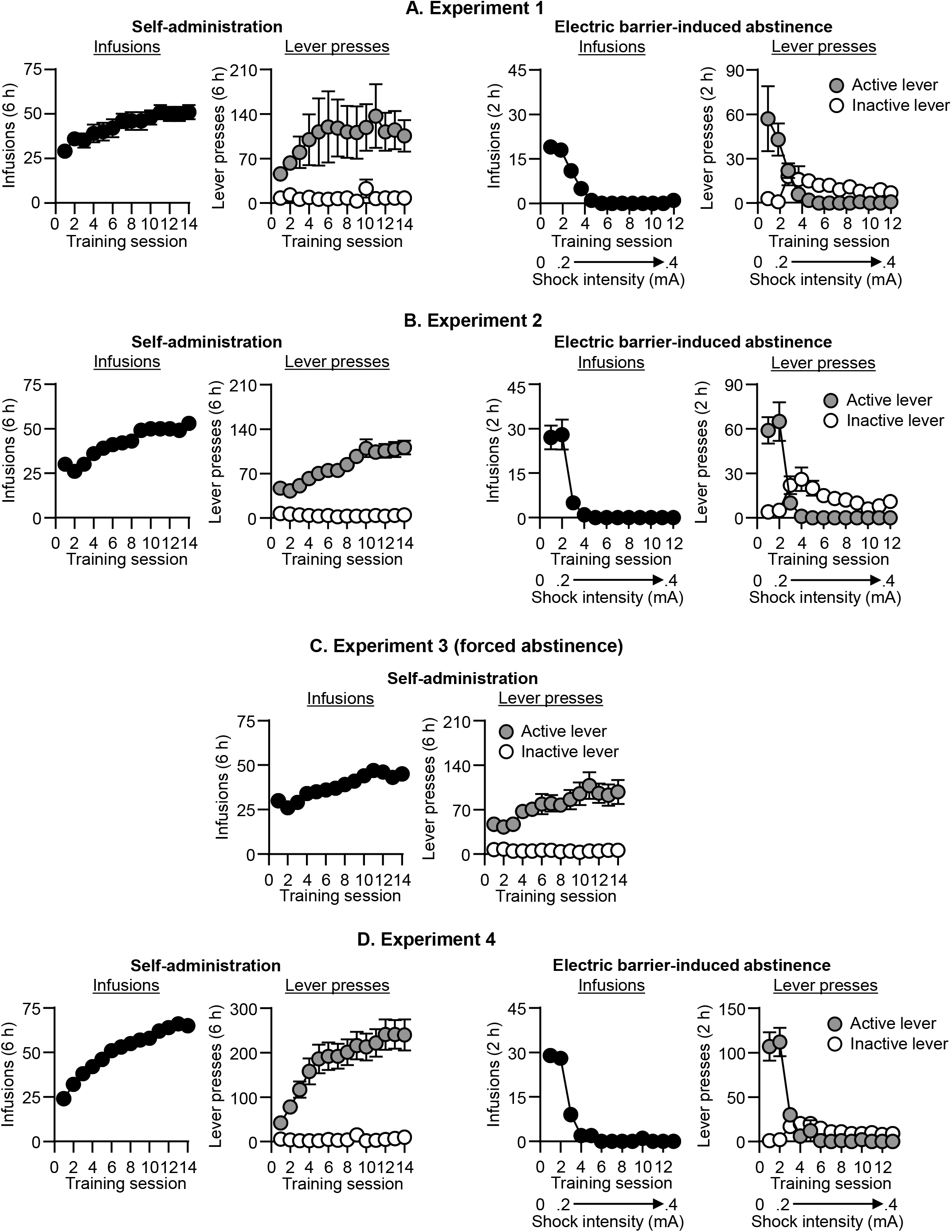
Oxycodone self-administration and electric barrier-induced abstinence (Exp. 1-4). <u>Left: Self-administration training.</u> Mean±SEM number of infusions, and active and inactive lever presses during the training phase in Experiments 1-4 (total n=13, 48, 34, and 59, respectively). <u>Right: Electric barrier-induced abstinence.</u> Mean±SEM number of infusions, and active and inactive lever presses during the electric barrier phase in Experiments 1-2, and 4.

### Electric barrier-induced abstinence

During this phase, oxycodone was available for 2 h per day for 13 or 16 days. We used the same parameters (oxycodone dose, reinforcement schedule, tone-light cues, etc.) that we used during the training phase. We achieved abstinence by introducing an electric barrier near the active lever (“shock zone”) (Cooper et al., 2007; Fredriksson et al., 2020). We separated the “shock zone” (2/3 of the chamber) from a “safe zone” (remaining 1/3 of the chamber) with a plastic demarcation (Mcmaster, cat# 9852K61). If the rats approached the active lever, they received a continuous mild footshock (0.1– 0.4 mA). On the first day, the current was set at 0.0 mA and current intensity was gradually increased to 0.3 mA by 0.1 mA increments per day. If the rats did not suppress oxycodone self-administration (< 3 infusions per day), we increased the shock intensity to 0.4 mA the next day. Prior to the electric barrier phase, we tested the rats’ sensitivity to footshock (operationally defined as the minimal shock level that causes the withdrawal of the front paw). There were no group or sex differences in shock sensitivity in any of the experiments, assessed with 0.05 mA increments, starting at 0.05 mA, and the values for individual rats ranged from 0.15 mA to 0.2 mA.

### Forced abstinence

During the forced abstinence phase, we moved the rats from the animal facility to their operant self-administration chamber and left them in the chamber for 2 h per day without turning on the electric barrier program.

### Relapse (incubation) tests

We tested the rats for oxycodone seeking for 30 or 90 min under extinction conditions during early (day 1) or late (day 15 or 18) abstinence, or both. During testing, we turned off the electric barrier and removed the plastic demarcation. We gave all rats a 30 min habituation period in the self-administration chamber before the start of the test session to allow them to realize that the barrier is not electrified. Lever presses resulted in delivery of the oxycodone-paired tone-light cue and activation of the infusion pump, but no drug infusions.

#### Exp. 1: vSub Fos expression after a test for incubated oxycodone seeking on day 15

The goal of Exp. 1 was to determine whether incubation of oxycodone seeking after electric barrier-induced abstinence is associated with increased Fos expression in vSub after the 90 min day 15 relapse test. The experiment consisted of four phases (Fig. 2A): Oxycodone self‐administration training (14 days), early tests for oxycodone seeking (abstinence day 1), electric barrier-induced abstinence (13 days), and late tests for oxycodone seeking (abstinence day 15). We used two groups of male and female rats (n=6-7 per group) in an experimental design that included the between-subjects factor of test condition (No-test, Test). We matched the groups for oxycodone intake during the training and electric barrier phases. On abstinence day 15, all rats went through a 90-min relapse test, and only rats from the test group were perfused immediately after the relapse test. We perfused rats from the no-test group the following day. To verify that incubation had occurred, we compared the number of lever presses during the 30 min day 1 test to the number of lever presses during the first 30 min of the day 15 test.

**Figure 2.**
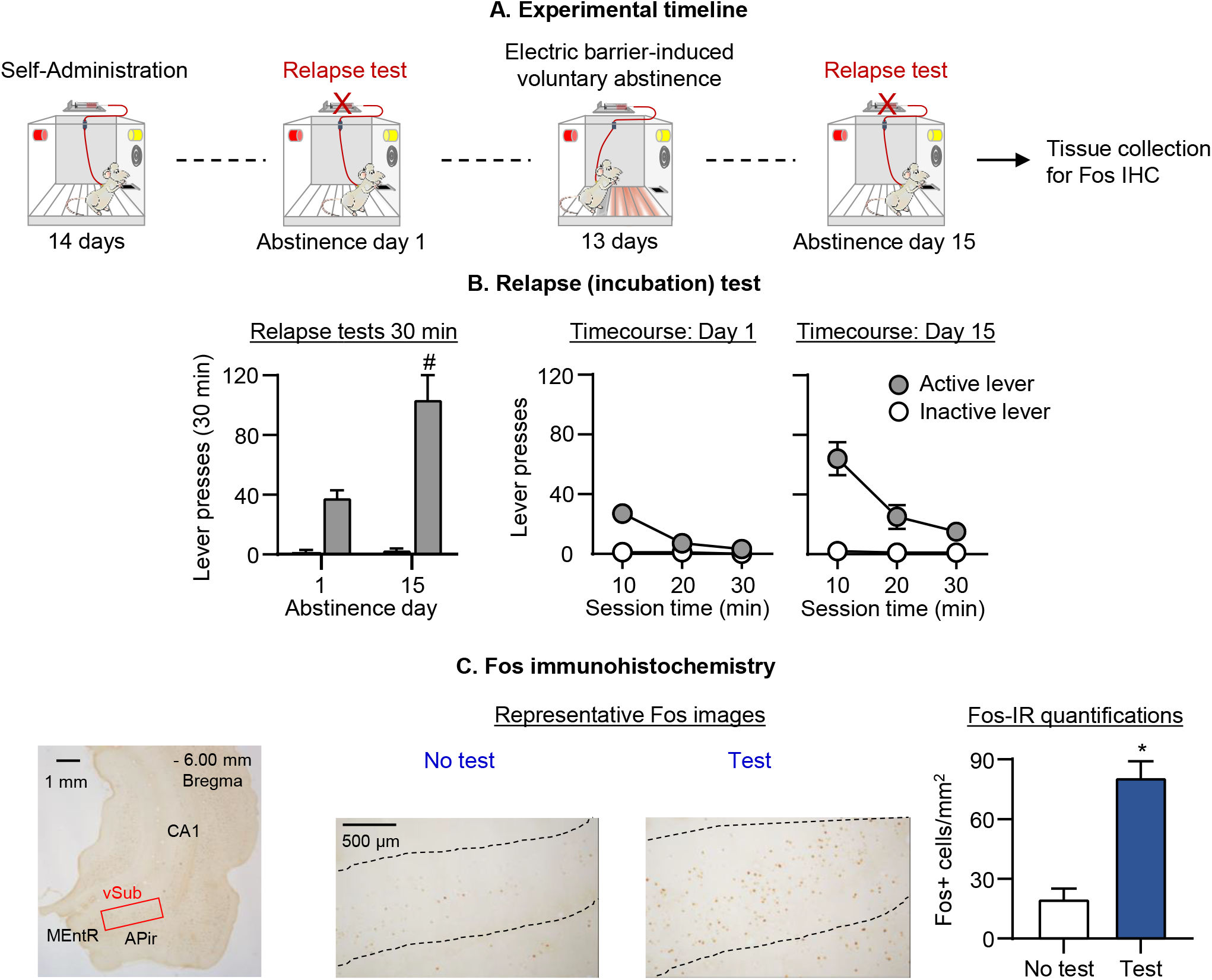
Effect of incubation of oxycodone seeking after electric barrier-induced abstinence on vSub Fos expression. (**A)** Timeline of Exp. 1. (**B**) <u>Relapse (incubation) tests:</u> Mean±SEM number of active lever presses during the 30-min day 1 test session and the first 30-min of the day 15 test session. During testing, active lever presses led to contingent presentation of the tone-light cue previously paired with oxycodone infusions during training, but not oxycodone infusions (extinction conditions). We tested the rats on days 1 and 15 (within-subjects design). (**C**) Representative images of Fos+ cells in vSub at 10X magnification and Fos-IR quantification: number of Fos+ cells (counts/mm^2^) in rats perfused either immediately after the day 15 test or 24-h later (No test, Test). * Different from No test. #Different from day 1, *p* < 0.05. Data are mean ± SEM. No test: n=6 (3 males, 3 females); Test: n=7 (4 males, 3 females). See Fig. S1 for individual data.

#### Exp. 2: Effect of muscimol-baclofen vSub inactivation on incubation after electric barrier-induced abstinence

In Exp. 1, we found that incubation of oxycodone seeking after electric barrier-induced abstinence was associated with increased Fos expression in vSub. Therefore, the goal of Exp. 2 was to determine whether vSub plays a causal role in incubation of oxycodone seeking after electric barrier-induced abstinence. For this purpose, we used the classical muscimol-baclofen inactivation procedure (McFarland and Kalivas, 2001). The experiment consisted of four phases (Fig. 3A): Oxycodone self‐administration training (14 days), early tests for oxycodone seeking (abstinence day 1), electric barrier-induced abstinence (13 days), and late tests for oxycodone seeking (abstinence day 15). We used 4 groups of male and female rats (n=10-14 per group) in a mixed experimental design that included the between-subjects factors of Abstinence day (1 or 15) and Muscimol-baclofen dose (0, 50+50 ng/side), and the within-subjects factor of Session time (30, 60, or 90 min). We matched the different groups for total oxycodone infusions during the training phase. We compared number of lever presses on day 1 and day 15 tests in rats injected with either saline or muscimol-baclofen into vSub, 30 min prior to the relapse tests on day 1 or day 15. We extracted brains after the tests to verify cannula placements.

**Figure 3.**
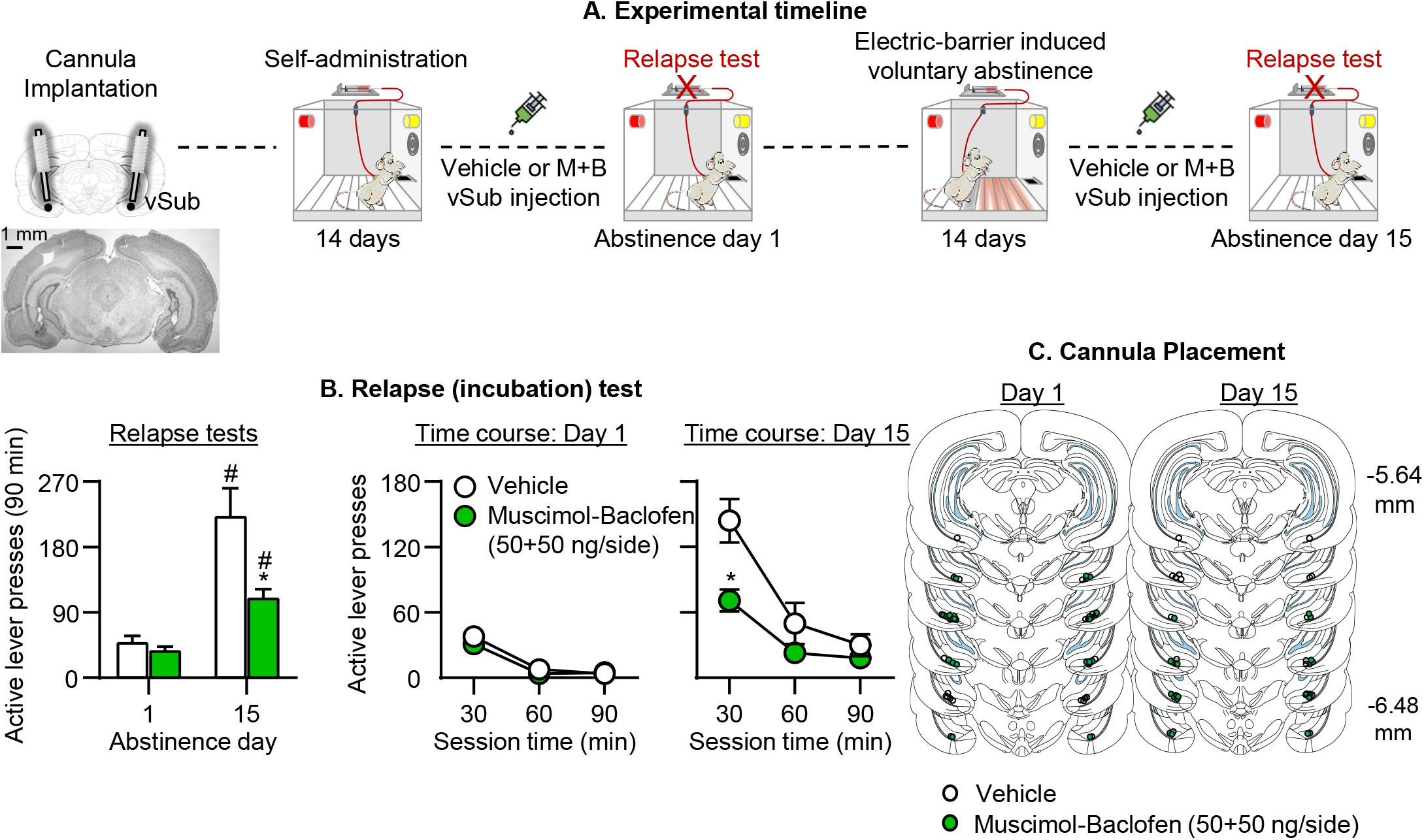
Effect of muscimol-baclofen vSub inactivation on incubation of oxycodone seeking after electric barrier-induced abstinence. (**A**) Timeline of Exp. 2. (**B**) <u>Relapse (incubation) tests:</u> Mean±SEM number of active lever presses during the 90-min test sessions after muscimol-baclofen vSub injection (0, 50+50 ng/side). We tested separate groups of rats on either day 1 or 15 (between-subjects design). (**C**) Cannula placements in vSub. * Different from vehicle. #Different from day 1, *p* < 0.05. Day 1: n=13-14 rats per dose (7-8 males/dose, 5-7 females/dose); Day 15: n=10-11 rats per dose (5 males/dose, 5-6 females/dose). See Fig. S1 for individual data.

#### Exp. 3: Effect of muscimol-baclofen vSub inactivation on incubation after forced abstinence

The goal of Exp. 3 was to test the specificity of the effect of muscimol-baclofen vSub inactivation on incubated oxycodone seeking after electric barrier-induced abstinence. For this purpose, we determined the effect of vSub vehicle (saline) or Muscimol-baclofen (50+50 ng/side) injections on incubated (day 15) oxycodone craving after forced abstinence. The experiment consisted of three phases (Fig. 4A): Oxycodone self‐administration training (14 days), forced abstinence (14 days), and late tests for oxycodone seeking (abstinence day 15). During the forced abstinence phase, we moved the rats from the animal facility to their operant self-administration chamber and left them in the chamber for 2 h per day without turning on the electric barrier program. We used 2 groups of male and female rats (n=16-18 per group) in a mixed experimental design that included the between-subjects factor of Muscimol-baclofen dose (0, 50+50 ng/side) and the within-subjects factor of Session time (30, 60, or 90 min). We matched the different groups for total oxycodone infusions during the training phase. We compared the number of lever presses on the day 15 relapse test in rats injected with either saline or muscimol-baclofen into the vSub 15 min prior to testing. We extracted the brains after the test to verify cannula placements.

**Figure 4.**
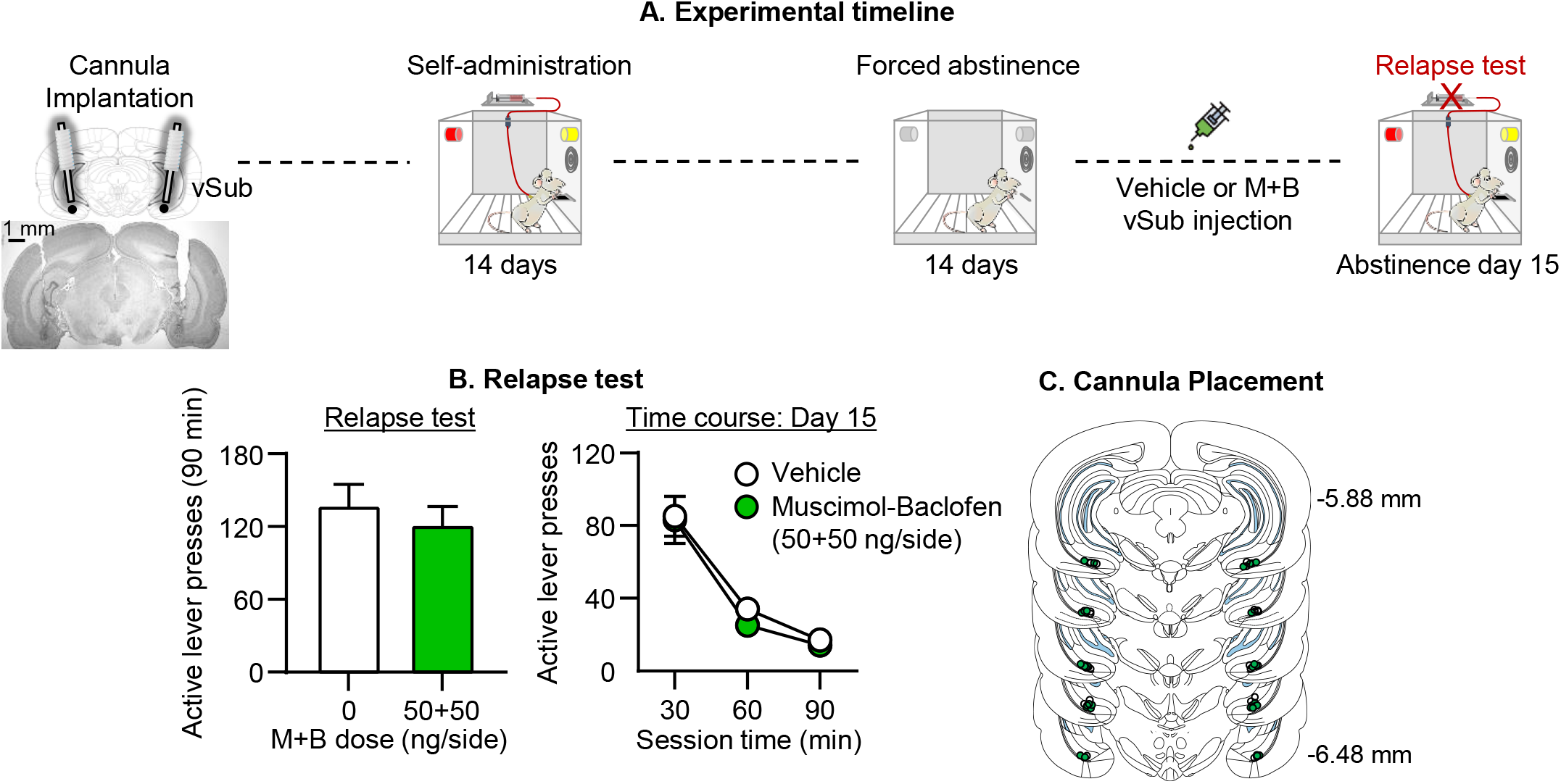
Effect of muscimol-baclofen vSub inactivation on incubation of oxycodone seeking after forced abstinence. (**A**) Timeline of Exp. 3. (**B**) <u>Relapse (incubation) test:</u> Mean±SEM number of active lever presses during the 90-min day 15 test session after muscimol-baclofen vSub injection (0, 50+50 ng/side). (**C**) Cannula placements. n=16-18 rats per dose (8-10 males/dose, 8 females/dose). See Fig. S1 for individual data.

#### Exp. 4: Effect of Daun02 vSub inactivation on incubation after electric barrier-induced abstinence

In Exp. 4, we used the Daun02 inactivation procedure (Koya et al., 2009) to determine whether activated neuronal ensembles in vSub play a causal role in incubated oxycodone seeking after electric barrier-induced abstinence. The experiment consisted of five phases (Fig. 5A): Oxycodone self‐administration training (14 days), electric barrier-induced abstinence (14 days), induction session (abstinence day 15), electric barrier induced abstinence (2 days) and late tests for oxycodone seeking (abstinence day 18). On the induction session (abstinence day 15), we briefly exposed groups of rats for 15 min to either the oxycodone self-administration context and cues associated with oxycodone injections (lever presses under extinction conditions) to induce relapse-dependent Fos in vSub, or a novel context (a plastic bowl with toys) to induce relapse-independent Fos in vSub. Next, 90 min after the induction sessions, when Fos and β-gal expression (Koya et al., 2009) are at their peak, we injected the rats with Daun02 (to inactivate the Fos-positive activated neurons) or vehicle. Between the induction day and the relapse test on abstinence 18, we conducted two additional electric barrier sessions. We used 4 groups of rats (n = 12-17 per group) and analyzed the data of the rats in each induction context (Novel context, Self-administration context) separately in a mixed experimental design with the between-subjects factor of Daun02 dose (0, 4 μg/side) and the within-subjects factor of Session time (30, 60, or 90 min). We matched the different groups for total oxycodone intake during the training and electric barrier phases. We compared number of active lever presses on the day 18 relapse test in rats injected with either saline or Daun02 dose (0, 4 µg/side) into the vSub on abstinence day 15. We perfused the rats after the test, verified cannula placements, and performed Fos and β-gal immunohistochemistry.

**Figure 5.**
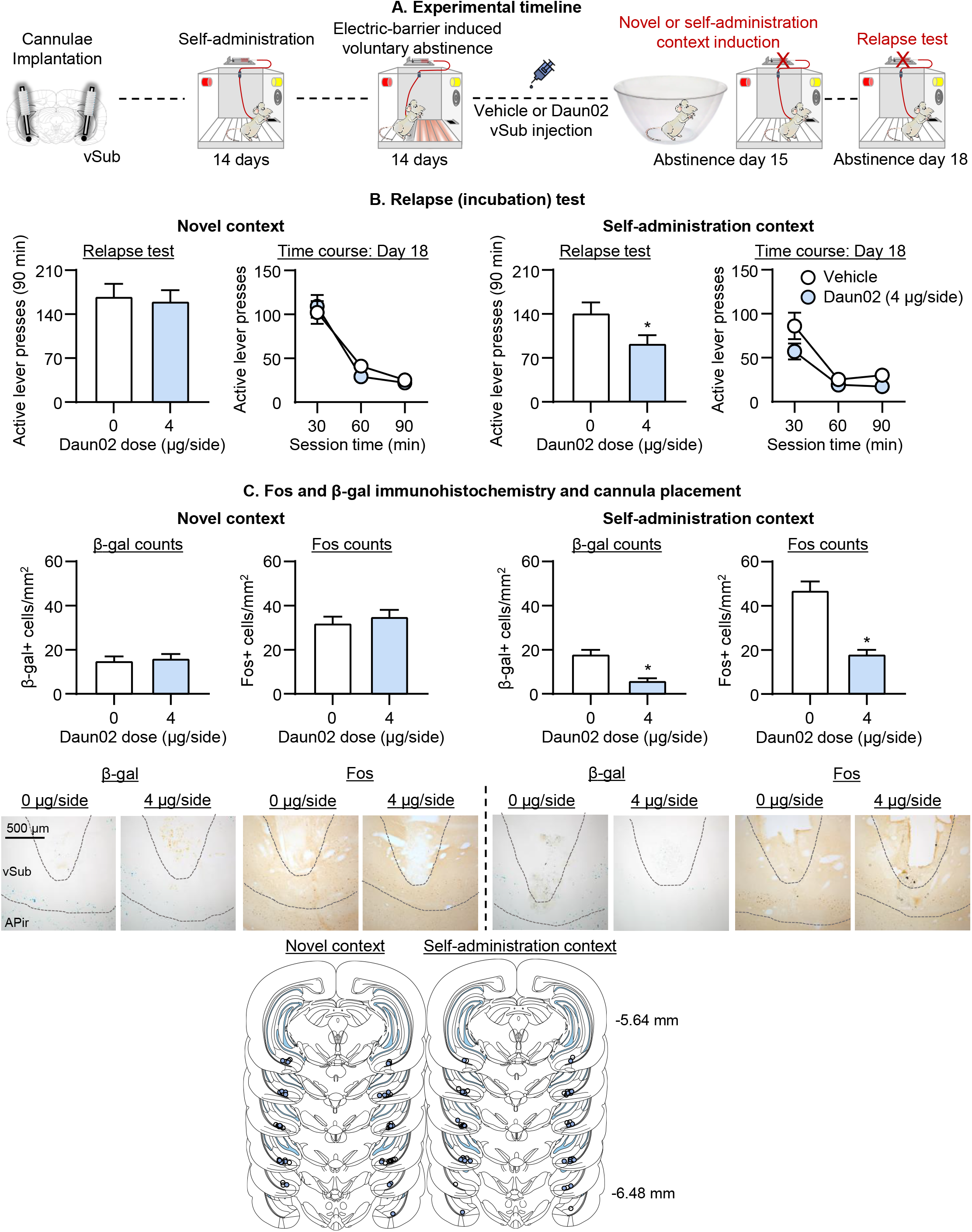
Effect of Daun02 vSub inactivation on incubation of oxycodone seeking after electric barrier-induced abstinence. (**A**) Timeline of Exp. 4. (**B**) <u>Relapse (incubation) tests:</u> Mean±SEM number of active lever presses during the day 18 90-min test session after Daun02 vSub injection (0, 4 µg/side) on day 15. (**C**) Representative images of β-gal+ and Fos+ cells in vSub at 10X magnification, β-gal and Fos-IR quantification: Mean±SEM number of β-gal+ or Fos+ cells (counts/mm^2^) in rats perfused after the day 18 relapse test, and cannula placements. * Different from vehicle, *p* < 0.05. Novel context: n=16-17 rats per dose (6-9 males/dose, 7-11 females/dose); Self-administration context: n=12-14 rats per dose (6-9 males/dose, 7-11 females/dose). See Fig. S1 for individual data.

### Statistical analyses

We analyzed the data with RM-ANOVAs, RM-ANCOVAs, ANOVAs, and ANCOVAs (inactive lever as a covariate) using SPSS (Version 24, GLM procedure). We followed significant main effects and interactions (p<0.05) with post-hoc tests (univariate ANOVAs or Fisher’s PLSD). We describe the different between- and within-subjects factors for the different analyses in the Results section. Because the multifactorial ANCOVAs yielded multiple main and interaction effects, we only report significant effects that are critical for data interpretation. For clarity, we indicate the results of post-hoc analyses with asterisks in the figures, but do not describe them in the Results section. For a complete reporting of the statistical analyses see Table S1, and for individual data of the bar graphs described in Fig. 2–5 see Fig. S1.

## Results

We used both male and female rats in all our experiments (Exp. 1-4). We did not use sex as a factor in the statistical analysis described below and in Table S1, because in our previous study, which was statistically powered to detect sex differences, we did not observe sex differences in oxycodone self-administration, electric barrier-induced abstinence, or incubation of oxycodone craving (Fredriksson et al., 2020).

### Oxycodone self-administration (Exp. 1-4)

The male and female rats demonstrated reliable oxycodone self-administration (Fig. 1A-D), as indicated by a significant increase in the number of infusions and active lever presses over the training days. The complete analyses for number of infusions and active and inactive lever presses during training are described in Table S1.

### Electric barrier-induced voluntary abstinence (Exp. 1, 2, and 4)

The male and female rats voluntarily abstained from drug self-administration when we introduced an electric barrier of increasing shock intensity near the active lever, as indicated by a significant decrease in the number of infusions and active lever presses during the abstinence phase (Fig.1A, B, and D). The mean number of infusions for the last 3 days of electric barrier-induced abstinence was less than 1 per session. In Exp. 1, Exp. 2, and Exp. 4, there were 7, 5, and 14 rats of the total rats (13, 21, and 59, respectively) in each experiment that required exposure to 0.4 mA to eliminate oxycodone self-administration. The statistical analyses of the electric barrier-induced abstinence phase are described in Table S1.

### Incubated oxycodone seeking after electric barrier-induced abstinence is associated with increased vSub activity

The goal of Exp. 1 was to determine whether incubation of oxycodone seeking after electric barrier-induced abstinence is associated with increased neuronal activity (assessed by Fos expression) in vSub. For this purpose, we tested male and female rats for oxycodone seeking under extinction conditions 1 day after oxycodone self-administration training and then tested them again after 15 days of electric barrier-induced abstinence. We perfused rats either immediately following the day 15 test (Test condition) or the following day (No-test condition).

#### Relapse (incubation) test

Oxycodone seeking in the relapse (incubation) tests was greater after 15 abstinence days than after 1 day, demonstrating incubation of oxycodone craving after electric barrier-induced abstinence (Fig. 2B). The two-way RM-ANCOVA (inactive lever as a covariate) for number of active lever presses of rats tested on day 1 and day 15, which included the within-subjects factors of Abstinence day (1 or 15), and Session time (30, 60, or 90 min), showed significant effects of Abstinence day (F_1,10_=6.2, p=0.032), Session time (F_2,20_=17.1, p<0.001), and Abstinence day x Session time (F_2,20_=7.0, p=0.005).

#### Fos immunohistochemistry

We measured Fos expression after the day 15 relapse (incubation) test. Relapse to oxycodone seeking during testing was associated with increased Fos expression in vSub (Fig. 2C). The one-way ANOVA for Fos+ cells/mm^2^in vSub, which included the between-subjects factor of Test condition (No-test, Test), showed a significant effect of this factor (F_1,11_=36.3, p<0.001). Representative pictures of Fos expression in the Test and No-test rats are shown in Fig. 2C.

The results of Exp. 1 demonstrate that incubation of oxycodone seeking after electric barrier-induced voluntary abstinence is associated with increased neuronal activity in vSub.

### Global vSub inactivation decreased incubated oxycodone seeking after electric barrier-induced abstinence

In Exp. 1 we found that incubation of oxycodone seeking after electric barrier-induced voluntary abstinence was associated with increased Fos expression in vSub. The goal of Exp. 2 was to determine a causal role of vSub in this form of incubation, using the classical muscimol-baclofen inactivation procedure (McFarland and Kalivas, 2001). We tested different groups of male and female rats for the effect of vSub saline or muscimol-baclofen injections on oxycodone seeking 1 day after oxycodone self-administration training or after 15 days of electric barrier-induced abstinence.

#### Relapse test

Muscimol-baclofen vSub inactivation decreased incubated oxycodone seeking on day 15 but had no effect on non-incubated oxycodone seeking on day 1 (Fig. 3B). The mixed factorial ANCOVA (inactive lever as a covariate) for number of active lever presses, which included the between-subjects factors of Abstinence day (1 or 15) and Muscimol-baclofen dose (0, 50+50 ng/side), and the within-subjects factor of Session time (30, 60, or 90 min), showed significant effects of Abstinence day (F_1,43_=43.3, p<0.001, Muscimol-baclofen dose (F_1,43_=10.5, p=0.002), Session time (F_2,86_=62.7, p<0.001), and Abstinence day x Muscimol-baclofen dose (F_1,43_=7.3, p=0.01).

The results of Exp. 2 demonstrate that global inhibition of vSub neuronal activity selectively decreased incubated, but not non-incubated, oxycodone seeking after electric barrier-induced voluntary abstinence.

### Global vSub inactivation had no effect on incubated oxycodone seeking after forced abstinence

The goal of Exp. 3 was to determine whether the effect of muscimol-baclofen vSub inactivation on incubated oxycodone seeking after electric barrier-induced voluntary abstinence would generalize to incubation after forced abstinence. For this purpose, we tested different groups of rats for the effect of saline or muscimol- baclofen injections into the vSub on oxycodone seeking after 15 days of forced abstinence from oxycodone self-administration.

#### Relapse test

Inactivation of the vSub with muscimol-baclofen had no effect on incubated oxycodone seeking on day 15 after forced abstinence (Fig. 4B). The mixed factorial ANCOVA (inactive lever as a covariate) for number of active lever presses of rats on day 15, which included the between-subjects factor of Muscimol-baclofen dose (0, 50+50 ng/side) and the within-subjects factor of Session time (30, 60, 90 min), showed a significant effect of Session time (F_2,62_=30.2, p<0.001) but no significant effects of Muscimol-baclofen dose or interaction (p values>0.1).

The results of Exp. 3 demonstrate that global inhibition of vSub neuronal activity had no effect on incubation of oxycodone seeking after forced abstinence.

### Selective inactivation of vSub neuronal ensembles decreased incubated oxycodone seeking after electric barrier-induced abstinence

In Exp. 2 we found that muscimol-baclofen inactivation of vSub decreased incubated oxycodone seeking after electric barrier-induced abstinence. These data indicate that vSub activity contributes to the incubated oxycodone seeking. However, from the perspective of the putative role of vSub activated neuronal ensembles and their role in incubation, the interpretation of the muscimol-baclofen data is confounded by the fact that this manipulation inhibits all neurons in a given brain area, independent of their task-related activity (incubated oxycodone seeking in our study). Therefore, the goal of Exp. 4 was to determine whether selective inactivation of oxycodone relapse-associated vSub neuronal ensembles would decrease incubation of oxycodone seeking after electric barrier-induced abstinence.

We trained male and female *Fos-LacZ* rats for 14 days of oxycodone self-administration and 14 days of electric barrier-induced abstinence. One day later (abstinence day 15), we exposed different groups of rats to either a 15 min “induction” session under extinction conditions in the self-administration chambers, during which lever presses were reinforced by the oxycodone-paired cues but not oxycodone (to activate the putative relapse-associated neuronal ensembles), or 15 min in a novel context (to induce relapse-independent activity). Ninety min after the induction session, we injected the rats with either saline or Daun02 (4 µg/side) into the vSub to inactivate the relapse-associated and relapse-independent Fos+ neurons, and tested all rats 3 days later (abstinence day 18) for incubated oxycodone seeking.

#### Induction day (day 15)

On induction day, active lever pressing in the 15 min oxycodone seeking session was not statistically different between the groups that received vehicle or Daun02 injections 90 min after session’s onset (mean±SEM active lever presses per 15 min: vehicle, 91±18; Daun02, 81±10; F_1,23_=0.02, p=0.90).

#### Relapse test (day 18)

We analyzed the data separately for saline vs. Daun02 for the two induction conditions (drug seeking session in the self-administration chambers versus novel context), because their experimental conditions prior to the relapse test were different (no oxycodone seeking under extinction conditions for the novel context group). Daun02 inactivation of vSub neurons activated during the short 15 min oxycodone seeking session (abstinence day 15) decreased oxycodone seeking 3 days later (Fig. 5B). The mixed factorial ANCOVA (inactive lever as a covariate) for number of active lever presses, which included the between-subjects factor of Daun02 dose (0, 4 µg/side) and the within-subjects factor of Session time (30, 60, or 90 min), showed significant effects of Daun02 dose (F_1,23_=4.5, p=0.044) and Session time (F_2,46_=20.9, p<0.001) but no significant interaction (p>0.1). In contrast, Daun02 inactivation of vSub neurons activated during a short 15 min novel context session (abstinence day 15) had no effect on oxycodone seeking 3 days later. The ANCOVA showed a significant effect of Session time (F_2,60_=39.1, p<0.01) but not Daun02 dose or an interaction (p values>0.1).

#### Fos and β-gal histochemistry

On the day 18 relapse test session, prior Daun02 injections decreased the number of both Fos and β-gal-labeled neurons in vSub of rats exposed to the short oxycodone-seeking session under extinction conditions on induction day but not in rats exposed to the novel context (Fig. 5C). These results indicate that the Daun02 manipulation selectively inactivated relapse-associated vSub neurons. One-way ANOVAs for Fos+ cells/mm^2^and β-gal+ cells/mm^2^in vSub, which included the between-subjects factor of Daun02 dose (0, 4 µg/side), showed a significant effect of this factor for Fos (F_1,24_=50.3, p<0.001) and β-gal (F_1,24_=33.1, p<0.001). In contrast, Daun02 inactivation of novel context-associated neurons in the vSub on induction day did not decrease Fos and β-gal-labeled neurons in vSub after the 90 min relapse test (p values>0.1). Representative pictures of Fos and β-gal staining are shown in Fig. 5C.

The results of Exp. 4 demonstrate a critical role of neuronal ensembles in the vSub for incubated oxycodone craving after electric barrier-induced voluntary abstinence.

## Discussion

We determined the role of vSub in incubation of opioid craving after voluntary abstinence induced by adverse consequences of opioid seeking. Incubated oxycodone seeking on abstinence day 15 was associated with increased vSub Fos expression. Additionally, global inactivation of vSub with muscimol-baclofen decreased incubated (day 15) but not non-incubated (day 1) oxycodone seeking. In contrast, muscimol-baclofen inactivation of vSub had no effect on incubated (day 15) oxycodone seeking after forced abstinence. Daun02 inactivation of vSub neurons previously activated by exposure to the oxycodone self-administration context and discrete cues during a short (15 min) “memory reactivation” session on induction day, decreased incubated oxycodone seeking and vSub neuronal activity 3 days later. In contrast, Daun02 inactivation of vSub neurons previously activated by novel context exposure on induction day had no effect. These results demonstrate a role of vSub activity in incubation of oxycodone seeking after voluntary abstinence, but not in incubation of oxycodone seeking after forced abstinence or non-incubated oxycodone seeking during early abstinence.

### Role of vSub in incubation of opioid craving after voluntary but not forced abstinence

Initial studies on the role of vSub or other ventral hippocampus areas in drug relapse/reinstatement have focused on psychostimulants. Electrical stimulation of vSub reinstates cocaine or amphetamine seeking after extinction (Vorel et al., 2001; Taepavarapruk et al., 2014). Muscimol-baclofen or lidocaine inactivation of ventral hippocampus or vSub decreases cue- and drug priming-induced reinstatement of cocaine and methamphetamine seeking (Sun and Rebec, 2003; Hiranita et al., 2006; Rogers and See, 2007). Muscimol-baclofen inactivation of ventral hippocampus decreases context-induced reinstatement of cocaine seeking (Lasseter et al., 2010). More recently, we showed a role of vSub and vSub→nucleus accumbens shell projections in context-induced reinstatement of heroin seeking after extinction (Bossert and Stern, 2014; Bossert et al., 2016) and context-induced relapse to alcohol seeking after punishment-induced abstinence (Marchant et al., 2016b). Together, the results from these studies indicate a general role of ventral hippocampus in relapse to drug seeking across drug classes and relapse models. However, the results of our study suggest that this general role does not generalize to incubation of drug seeking. As described above, our results demonstrate a selective role of vSub in incubated oxycodone seeking after electric barrier-induced abstinence but not incubated oxycodone seeking after forced abstinence or non-incubated oxycodone seeking during early abstinence.

The different effects of vSub inactivation on incubation of oxycodone seeking after electric barrier-induced voluntary abstinence versus forced abstinence are likely due to the different methods used to achieve abstinence (Marchant et al., 2019; Fredriksson et al., 2021). For example, basolateral amygdala activity plays opposite roles in context-induced relapse of cocaine seeking after extinction versus punishment (Pelloux et al., 2018). Additionally, PKCdelta and somatostatin in central amygdala play dissociable roles in incubation of methamphetamine craving after homecage forced abstinence versus prevention of this incubation after voluntary abstinence induced by rewarding social interaction (Venniro et al., 2020). Furthermore, at the behavioral level, incubation of oxycodone craving is strongly potentiated after electric barrier-induced voluntary abstinence compared to forced abstinence (Fredriksson et al., 2020). We speculate that one reason for this potentiation is stress exposure during the electric-barrier phase. A recent study, showing that repeated restraint stress exposure during forced abstinence, increases incubation of cocaine craving (Glynn et al., 2018) supports this speculation. Also, potentially relevant to our results is that ventral hippocampus inactivation with muscimol-baclofen decreases the potentiation effect of repeated restraint stress on resumption of nicotine self-administration when given during 8 abstinence days (Yu and Sharp, 2015).

### Role of vSub neuronal ensembles in incubation of opioid craving

We used the Daun02 inactivation method (Koya et al., 2009) to study the causal role of vSub neuronal ensembles in incubation of opioid craving after electric barrier-induced abstinence. This method is used to selectively inactivate behaviorally activated neurons (Fos-positive) and was developed to study the causal role of activated neuronal ensembles in conditioned drug effects, drug relapse, and learned behaviors (Koya et al., 2009; Cruz et al., 2013; Cruz et al., 2015). In this regard, muscimol-baclofen and other classical reversible inactivation methods, site-specific injections of receptor antagonists/agonists, permanent lesion methods, or more modern optogenetic and chemogenetic methods to inhibit specific cell-types are not suitable to study causal roles of neuronal ensembles in learned behaviors (Cruz et al., 2013). This is because these methods invariably inhibit both behaviorally activated and non-activated neurons within a particular brain area.

We found that vSub Daun02 inactivation decreased incubated oxycodone seeking after electric barrier-induced abstinence, demonstrating a role of vSub neuronal ensembles in this incubation. A question for future studies is which specific vSub cell types contribute to incubation of oxycodone craving. We speculate that the putative vSub incubation-related ensembles are not cell-type specific. In our previous relapse/reinstatement and incubation studies, we found no evidence for cell-type specificity (Drd1-versus Drd2-expressing cells in striatum or GABA- versus glutamate-expressing cells in mPFC) of Fos-expressing neuronal ensembles (Bossert et al., 2011; Fanous et al., 2012; Cruz et al., 2014; Warren et al., 2016; Caprioli et al., 2017).

### Methodological considerations

One potential issue is that the effects of muscimol-baclofen or Daun02 vSub injections on incubated oxycodone seeking are due to nonspecific performance deficits. This is unlikely because muscimol-baclofen vSub injections had no effect on incubated oxycodone seeking after forced abstinence. Additionally, vSub Daun02 injections after novel context exposure had no effect on incubated oxycodone seeking after electric barrier-induced abstinence.

Another issue is that Daun02 injections on induction day decreased incubated oxycodone seeking during testing 3 days later by nonspecifically inactivating a random group of Fos-positive neurons. This is unlikely because Daun02 inactivation of vSub neurons decreased incubated oxycodone seeking and vSub neuronal activity only when Daun02 was injected after exposure to the oxycodone self-administration context and discrete cues during a short “memory reactivation” induction session but not when Daun02 was injected after exposure to the novel context, a manipulation that increases Fos expression in vSub (Marchant et al., 2016a) and other cortical areas (Badiani et al., 1998). These results suggest a selective effect of Daun02 inactivation on incubation-associated neuronal ensembles and agree with those from studies showing that Daun02 inactivation inhibits incubation-related ensembles in other brain regions (Fanous et al., 2012; Funk et al., 2016; Caprioli et al., 2017).

Another issue is that the magnitude of the inhibitory effect of Daun02 inactivation on incubated oxycodone seeking (~35%, Fig. 5B) was weaker than that of muscimol+baclofen (~50%, Fig. 3B). In this regard, Daun02 inactivation only partially interferes with the function of incubation-related neuronal ensembles because in vSub (and other brain areas, see (Bossert et al., 2011; Fanous et al., 2012; Funk et al., 2016; Caprioli et al., 2017), β-gal is expressed in the majority (~75%, unpublished data) but not all Fos-expressing neurons. Additionally, incubated oxycodone seeking is likely also controlled by Fos-negative neurons not strongly activated on induction day and consequently not inactivated by Daun02.

A final methodological issue is that the number of Fos-positive neurons after the relapse test in the vehicle condition of the Daun02 experiment (Fig. 5C) was lower than the number of Fos-positive neurons under similar test conditions in Exp. 1 (Fig. 2C). Typically, there is a lower number of Fos-expressing neurons in intact brain tissue versus brain tissue collected after intracranial drug injections due to tissue damage (Caprioli et al., 2017; Venniro et al., 2017). In the same experiment, the number of β-gal-positive neurons was lower than the Fos-positive neurons (Fig. 5C, for similar pattern of results see (Koya et al., 2009; Bossert et al., 2011; Warren et al., 2016; Caprioli et al., 2017)). This difference is likely due to a difference in sensitivity of the X-gal assay (enzymatic assay that does not depend on an antibody binding to an antigen) versus Fos immunohistochemistry.

### Concluding remarks

We recently developed a rat model of incubation of craving after electric barrier-induced voluntary abstinence (Fredriksson et al., 2020; Fredriksson et al., 2021) and proposed that our model mimics, to some degree, human voluntary abstinence due to negative consequences of *drug seeking* prior to *drug taking* (Marlatt, 1996; Epstein and Preston, 2003). Here, we used the activity marker Fos and pharmacological and chemogenetic methods to demonstrate that vSub neuronal ensembles, which encode the learned associations between oxycodone reward and cues and contexts associated with oxycodone self-administration, contribute to incubation of opioid craving after abstinence induced by negative consequences of drug seeking. We also showed that vSub activity did not contribute to incubation of opioid craving after forced abstinence, suggesting distinct mechanisms of incubation after forced versus voluntary abstinence.

## Acknowledgments

The research was supported by funds from the Intramural Research Program of the NIDA-NIH (YS and BTH) and The Swedish Research Council International Postdoc grant (2019-00658) to IF.

## References

Badiani A, Oates MM, Day HE, Watson SJ, Akil H, Robinson TE (1998) Amphetamine-induced behavior, dopamine release, and c-fos mRNA expression: modulation by environmental novelty. J Neurosci 18:10579–10593.

Blackwood CA, Leary M, Salisbury A, McCoy MT, Cadet JL (2019) Escalated oxycodone self-administration causes differential striatal mRNA expression of FGFs and IEGs following abstinence-associated incubation of oxycodone craving. Neuroscience 415:173–183.

Bossert JM, Stern AL (2014) Role of ventral subiculum in context-induced reinstatement of heroin seeking in rats. Addict Biol 19:338–342.

Bossert JM, Stern AL, Theberge FR, Cifani C, Koya E, Hope BT, Shaham Y (2011) Ventral medial prefrontal cortex neuronal ensembles mediate context-induced relapse to heroin. Nat Neurosci 14:420–422.

Bossert JM, Stern AL, Theberge FRM, Marchant NJ, Wang H, Morales M, Shaham Y (2012) Role of projections from ventral medial prefrontal cortex to nucleus accumbens shell in context-induced reinstatement of heroin seeking. J Neurosci 32:4982–4991.

Bossert JM, Adhikary S, St Laurent R, Marchant NJ, Wang HL, Morales M, Shaham Y (2016) Role of projections from ventral subiculum to nucleus accumbens shell in context-induced reinstatement of heroin seeking in rats. Psychopharmacology 233:1991–2004.

Bossert JM, Hoots JK, Fredriksson I, Adhikary S, Zhang M, Venniro M, Shaham Y (2019) Role of mu, but not delta or kappa, opioid receptors in context-induced reinstatement of oxycodone seeking. Eur J Neurosci 50:2075–2085.

Caprioli D, Venniro M, Zhang M, Bossert JM, Warren BL, Hope BT, Shaham Y (2017) Role of dorsomedial striatum neuronal ensembles in incubation of methamphetamine craving after voluntary abstinence. J Neurosci 37:1014–1027.

Caprioli D, Venniro M, Zeric T, Li X, Adhikary S, Madangopal R, Marchant NJ, Lucantonio F, Schoenbaum G, Bossert JM, Shaham Y (2015) Effect of the novel positive allosteric modulator of metabotropic glutamate receptor 2 AZD8529 on incubation of methamphetamine craving after prolonged voluntary abstinence in a rat model. Biol Psychiatry 78:463–473.

Cooper A, Barnea-Ygael N, Levy D, Shaham Y, Zangen A (2007) A conflict rat model of cue-induced relapse to cocaine seeking. Psychopharmacology 194:117–125.

Cruz FC, Javier Rubio F, Hope BT (2015) Using c-fos to study neuronal ensembles in corticostriatal circuitry of addiction. Brain Res 1628:157–173.

Cruz FC, Koya E, Guez-Barber DH, Bossert JM, Lupica CR, Shaham Y, Hope BT (2013) New technologies for examining the role of neuronal ensembles in drug addiction and fear. Nat Rev Neurosci 14:743–754.

Cruz FC, Babin KR, Leao RM, Goldart EM, Bossert JM, Shaham Y, Hope BT (2014) Role of nucleus accumbens shell neuronal ensembles in context-induced reinstatement of cocaine-seeking. J Neurosci 34:7437–7446.

de Guglielmo G, Crawford E, Kim S, Vendruscolo LF, Hope BT, Brennan M, Cole M, Koob GF, George O (2016) Recruitment of a Neuronal Ensemble in the Central Nucleus of the Amygdala Is Required for Alcohol Dependence. J Neurosci 36:9446–9453.

Dong Y, Taylor JR, Wolf ME, Shaham Y (2017) Circuit and synaptic plasticity mechanisms of drug relapse. J Neurosci 37:10867–10876.

Engeln M, Bastide MF, Toulme E, Dehay B, Bourdenx M, Doudnikoff E, Li Q, Gross CE, Boue-Grabot E, Pisani A, Bezard E, Fernagut PO (2016) Selective Inactivation of Striatal FosB/DeltaFosB-Expressing Neurons Alleviates L-DOPA-Induced Dyskinesia. Biol Psychiatry 79:354–361.

Epstein DH, Preston KL (2003) The reinstatement model and relapse prevention: a clinical perspective. Psychopharmacology 168:31–41.

Fanous S, Goldart EM, Theberge FR, Bossert JM, Shaham Y, Hope BT (2012) Role of orbitofrontal cortex neuronal ensembles in the expression of incubation of heroin craving. J Neurosci 32:11600–11609.

Fredriksson I, Applebey SV, Minier-Toribio A, Shekara A, Bossert JM, Shaham Y (2020) Effect of the dopamine stabilizer (-)-OSU6162 on potentiated incubation of opioid craving after electric barrier-induced voluntary abstinence. Neuropsychopharmacology 45:770–779.

Fredriksson I, Venniro M, Reiner DJ, Chow JJ, Bossert JM, Shaham Y (2021) Animal models of drug relapse and craving after voluntary abstinence: a review. Pharmacol Rev (accepted pending revisions).

Funk D, Coen K, Tamadon S, Hope BT, Shaham Y, Lê AD (2016) Role of central amygdala neuronal ensembles in incubation of nicotine craving. J Neurosci 36(33):8612–8623.

Glynn RM, Rosenkranz JA, Wolf ME, Caccamise A, Shroff F, Smith AB, Loweth JA (2018) Repeated restraint stress exposure during early withdrawal accelerates incubation of cue-induced cocaine craving. Addict Biol 23:80–89.

Grimm JW, Hope BT, Wise RA, Shaham Y (2001) Incubation of cocaine craving after withdrawal. Nature 412:141–142.

Hiranita T, Nawata Y, Sakimura K, Anggadiredja K, Yamamoto T (2006) Suppression of methamphetamine-seeking behavior by nicotinic agonists. Proc Natl Acad Sci U S A 103:8523–8527.

Ito R, Lee ACH (2016) The role of the hippocampus in approach-avoidance conflict decision-making: Evidence from rodent and human studies. Behav Brain Res 313:345–357.

Kane L, Venniro M, Quintana-Feliciano R, Madangopal R, Rubio FJ, Bossert JM, Caprioli D, Shaham Y, Hope BT, Warren BL (2020) Fos-expressing neuronal ensemble in rat ventromedial prefrontal cortex encodes cocaine seeking but not food seeking in rats. Addict Biol:e12943.

Kasof GM, Smeyne RJ, Curran T, Morgan JI (1996) Developmental expression of Fos-lacZ in the brains of postnatal transgenic rats. Brain Res Dev Brain Res 93:191–197.

Koya E, Golden SA, Harvey BK, Guez-Barber DH, Berkow A, Simmons DE, Bossert JM, Nair SG, Uejima JL, Marin MT, Mitchell TB, Farquhar D, Ghosh SC, Mattson BJ, Hope BT (2009) Targeted disruption of cocaine-activated nucleus accumbens neurons prevents context-specific sensitization. Nat Neurosci 12:1069–1073.

Laque A, G Ldn, Wagner GE, Nedelescu H, Carroll A, Watry D, T Mk, Koya E, Hope BT, Weiss F, Elmer GI, Suto N (2019) Anti-relapse neurons in the infralimbic cortex of rats drive relapse-suppression by drug omission cues. Nat Commun 10:3934.

Lasseter HC, Xie X, Ramirez DR, Fuchs RA (2010) Sub-region specific contribution of the ventral hippocampus to drug context-induced reinstatement of cocaine-seeking behavior in rats. Neuroscience 171:830–839.

Marchant NJ, Campbell EJ, Pelloux Y, Bossert JM, Shaham Y (2019) Context-induced relapse after extinction versus punishment: similarities and differences. Psychopharmacology 236:439–448.

Marchant NJ, Whitaker LR, Bossert JM, Harvey BK, Hope BT, Kaganovsky K, Adhikary S, Prisinzano TE, Vardy E, Roth BL, Shaham Y (2016a) Behavioral and physiological effects of a novel kappa-opioid receptor-based DREADD in Rats. Neuropsychopharmacology 41:402–409.

Marchant NJ, Campbell EJ, Whitaker LR, Harvey BK, Kaganovsky K, Adhikary S, Hope BT, Heins RC, Prisinzano TE, Vardy E, Bonci A, Bossert JM, Shaham Y (2016b) Role of ventral subiculum in context-induced relapse to alcohol seeking after punishment-imposed abstinence. J Neurosci 36:3281–3294.

Marlatt AG (1996) Models of relapse and relapse prevention: a commentary. Exp Clin Psychopharmacol 4:55–60.

McFarland K, Kalivas PW (2001) The circuitry mediating cocaine-induced reinstatement of drug-seeking behavior. J Neurosci 21:8655–8663.

Morgan JI, Curran T (1991) Stimulus-transcription coupling in the nervous system: involvement of the inducible proto-oncogenes fos and jun. Annu Rev Neurosci 14:421–451.

Nunes EV, Gordon M, Friedmann PD, Fishman MJ, Lee JD, Chen DT, Hu MC, Boney TY, Wilson D, O’Brien CP (2018) Relapse to opioid use disorder after inpatient treatment: Protective effect of injection naltrexone. J Subst Abuse Treat 85:49–55.

O’Brien CP, Childress AR, McLellan AT, Ehrman R (1992) Classical conditioning in drug-dependent humans. Ann N Y Acad Sci 654:400–415.

Pelloux Y, Minier-Toribio A, Hoots JK, Bossert JM, Shaham Y (2018) Opposite effects of basolateral amygdala inactivation on context-induced relapse to cocaine seeking after extinction versus punishment. J Neurosci 38:51–59.

Pfarr S, Meinhardt MW, Klee ML, Hansson AC, Vengeliene V, Schonig K, Bartsch D, Hope BT, Spanagel R, Sommer WH (2015) Losing Control: Excessive Alcohol Seeking after Selective Inactivation of Cue-Responsive Neurons in the Infralimbic Cortex. J Neurosci 35:10750–11061.

Pickens CL, Airavaara M, Theberge F, Fanous S, Hope BT, Shaham Y (2011) Neurobiology of the incubation of drug craving. Trends Neurosci 34:411–420.

Reiner DJ, Fredriksson I, Lofaro OM, Bossert JM, Shaham Y (2019) Relapse to opioid seeking in rat models: behavior, pharmacology and circuits. Neuropsychopharmacology 44:465–477.

Reiner DJ, Lofaro OM, Applebey SV, Korah H, Venniro M, Cifani C, Bossert JM, Shaham Y (2020) Role of projections between piriform cortex and orbitofrontal cortex in relapse to fentanyl seeking after palatable food choice-induced voluntary abstinence. J Neurosci 40:2485–2497.

Rogers JL, See RE (2007) Selective inactivation of the ventral hippocampus attenuates cue-induced and cocaine-primed reinstatement of drug-seeking in rats. Neurobiol Learn Mem 87:688–692.

Rudd RA, Seth P, David F, Scholl L (2016) Increases in drug and opioid-involved overdose deaths - United States, 2010-2015. MMWR Morb Mortal Wkly Rep 65:1445–1452.

Schumacher A, Villaruel FR, Ussling A, Riaz S, Lee ACH, Ito R (2018) Ventral Hippocampal CA1 and CA3 Differentially Mediate Learned Approach-Avoidance Conflict Processing. Curr Biol 28:1318–1324 e1314.

Shalev U, Morales M, Hope B, Yap J, Shaham Y (2001) Time-dependent changes in extinction behavior and stress-induced reinstatement of drug seeking following withdrawal from heroin in rats. Psychopharmacology 156:98–107.

Sinha R (2011) New findings on biological factors predicting addiction relapse vulnerability. Curr Psychiatry Rep 13:398–405.

Stopper CM, Floresco SB (2014) What’s better for me? Fundamental role for lateral habenula in promoting subjective decision biases. Nat Neurosci 17:33–35.

Sun W, Rebec GV (2003) Lidocaine inactivation of ventral subiculum attenuates cocaine-seeking behavior in rats. J Neurosci 23:10258–11064.

Taepavarapruk P, Butts KA, Phillips AG (2014) Dopamine and glutamate interaction mediates reinstatement of drug-seeking behavior by stimulation of the ventral subiculum. Int J Neuropsychopharmacol 18.

Theberge FR, Pickens CL, Goldart E, Fanous S, Hope BT, Liu QR, Shaham Y (2012) Association of time-dependent changes in mu opioid receptor mRNA, but not BDNF, TrkB, or MeCP2 mRNA and protein expression in the rat nucleus accumbens with incubation of heroin craving. Psychopharmacology 224:559–571.

Venniro M, Caprioli D, Shaham Y (2016) Animal models of drug relapse and craving: From drug priming-induced reinstatement to incubation of craving after voluntary abstinence. Prog Brain Res 224:25–52.

Venniro M, Russell TI, Ramsey LA, Richie CT, Lesscher HM, Giovanetti SM, Messing RO, Shaham Y (2020) Abstinence-dependent dissociable central amygdala microcircuits control drug craving. Proceedings of the National Academy of Sciences 117:8126–8134.

Venniro M, Zhang M, Caprioli D, Hoots JK, Golden SA, Heins C, Morales M, Epstein DH, Shaham Y (2018) Volitional social interaction prevents drug addiction in rat models. Nat Neurosci 21:1520–1529.

Venniro M, Caprioli D, Zhang M, Whitaker LR, Zhang S, Warren BL, Cifani C, Marchant NJ, Yizhar O, Bossert JM, Chiamulera C, Morales M, Shaham Y (2017) The anterior insular cortex-->central amygdala glutamatergic pathway Is critical to relapse after contingency management. Neuron 96:414–427 e418.

Vorel SR, Liu X, Hayes RJ, Spector JA, Gardner EL (2001) Relapse to cocaine-seeking after hippocampal theta burst stimulation. Science 292:1175–1178.

Warren BL, Mendoza MP, Cruz FC, Leao RM, Caprioli D, Rubio FJ, Whitaker LR, McPherson KB, Bossert JM, Shaham Y, Hope BT (2016) Distinct Fos-expressing neuronal ensembles in the ventromedial prefrontal cortex mediate food reward and extinction memories. J Neurosci 36:6691–6703.

Warren BL, Kane L, Venniro M, Selvam P, Quintana-Feliciano R, Mendoza MP, Madangopal R, Komer L, Whitaker LR, Rubio FJ, Bossert JM, Caprioli D, Shaham Y, Hope BT (2019) Separate vmPFC ensembles control cocaine self-administration versus extinction in rats. J Neurosci 39:7394–7407.

Wikler A (1973) Dynamics of drug dependence. Implications of a conditioning theory for research and treatment. Arch Gen Psychiatry 28:611–616.

Wolf ME (2016) Synaptic mechanisms underlying persistent cocaine craving. Nat Rev Neurosci 17:351–365.

Yu G, Sharp BM (2015) Basolateral amygdala and ventral hippocampus in stress-induced amplification of nicotine self-administration during reacquisition in rat. Psychopharmacology (Berl) 232:2741–2749.

