## Supplemental information for "Role of ventral subiculum neuronal ensembles in incubation of oxycodone craving after electric barrier-induced voluntary abstinence"

Table of content

Table S1. Statistical results
Figure S1. Individual data

**Supplementary Table 1**. Statistical analysis for the behavioral results (SPSS GLM repeated-measures module). Partial Eta$^2$ = proportion of explained variance. For ANCOVAs, the covariate is inactive lever presses. RM, repeated measures; SA, self-administration; M+B, muscimol-baclofen; vSub, ventral subiculum

**Exp. 1.** vSub Fos expression after a test for incubated oxycodone craving on day 15

| Figure number | Factor name | F-value | *p*-value | Partial Eta$^2$ |
|---|---|---|---|---|
| Figure 1B. Self-administration training<br>Infusions<br>RM-ANOVA<br>Within-subjects factor: Session | Session (1-14) within-subjects | $F_{13,156}$ = 10.5 | <0.001* | 0.466 |
| Figure 1B. Self-administration training<br>Active lever presses<br>RM-ANOVA<br>Within-subjects factors: Session, Lever | Session (1-14) within-subjects<br>Lever (Active, Inactive) within subjects<br>Lever X Session interaction | $F_{13,156}$ = 1.8<br>$F_{1,12}$ = 7.7<br>$F_{13,156}$ = 1.9 | 0.053<br>0.017*<br>0.030* | 0.128<br>0.390<br>0.139 |
| Figure 1C. Electric barrier<br>Infusions<br>RM-ANOVA<br>Within-subjects factor: Session | Session (1-13) within-subjects | $F_{12,144}$ = 49.3 | <0.001* | 0.804 |
| Figure 1C. Electric barrier<br>Active lever presses<br>RM-ANOVA<br>Within-subjects factors: Session, Lever | Session (1-13) within-subjects<br>Lever (Active, Inactive) within subjects<br>Lever X Session interaction | $F_{12,144}$ = 5.4<br>$F_{1,12}$ = 0.02<br>$F_{12,144}$ = 8.0 | <0.001*<br>0.898<br><0.001* | 0.309<br>0.001<br>0.401 |
| Figure 1D. Relapse tests days 1 and 15 (Total 30 min)<br>Active lever presses<br>RM-ANCOVA<br>Within-subjects factor: Abstinence day<br>Covariates: Inactive day 1, Inactive day 15 (30 min) | Abstinence day (1,15) within-subjects | $F_{1,10}$ = 6.2 | 0.032* | 0.384 |

| | | | | |
|---|---|---|---|---|
| Figure 1D. Relapse tests days 1 and 15 (30 min time course)<br><u>Active lever presses</u><br>RM-ANCOVA<br>Within-subjects factors: Abstinence day, Session time<br>Covariates: Inactive day 1, Inactive day 15 (30 min) | Abstinence day (1,15) within-subjects<br>Session time (10, 20, 30) within-subjects<br>Abstinence day X Session time interaction | $F_{1,10}$ = 6.2<br>$F_{2,20}$ = 17.1<br>$F_{2,20}$ = 7.0 | 0.032*<br><0.001*<br>0.005* | 0.384<br>0.631<br>0.412 |
| Figure 1E. Fos immunohistochemistry<br><u>vSub Fos Counts</u><br>One-way ANOVA<br>Between-subjects factor: Test condition | Test condition (No Test, Test) between-subjects | $F_{1,11}$ = 36.3 | <0.001* | 0.768 |

**Exp. 2.** Effect of muscimol-baclofen vSub inactivation on incubation after electric barrier-induced abstinence

| Figure number | Factor name | F-value | *p*-value | Partial Eta$^2$ |
|---|---|---|---|---|
| Figure 2B. Self-administration training<br><u>Infusions</u><br>RM-ANOVA<br>Within-subjects factor: Session | Session (1-14) within-subjects | $F_{13,611}$ = 41.2 | <0.001* | 0.467 |
| Figure 2B. Self-administration training<br><u>Active lever presses</u><br>RM-ANOVA<br>Within-subjects factors: Session, Lever | Session (1-14) within-subjects<br>Lever (Active, Inactive) within subjects<br>Lever X Session interaction | $F_{13,611}$ = 16.3<br>$F_{1,47}$ = 134.6<br>$F_{13,611}$ = 17.9 | <0.001*<br><0.001*<br><0.001* | 0.257<br>0.741<br>0.276 |
| Figure 2C. Electric barrier<br><u>Infusions</u><br>RM-ANOVA<br>Within-subjects factor: Session | Session (1-12) within-subjects | $F_{11,220}$= 30.5 | <0.001* | 0.604 |
| Figure 2C. Electric barrier<br><u>Active lever presses</u><br>RM-ANOVA<br>Within-subjects factors: Session, Lever | Session (1-12) within-subjects<br>Lever (Active, Inactive) within subjects<br>Lever X Session interaction | $F_{11,220}$ = 15.9<br>$F_{1,20}$ = 0.1<br>$F_{11,220}$ = 26.5 | <0.001*<br>0.73<br><0.001* | 0.442<br>0.006<br>0.57 |
| Figure 2D. Relapse tests days 1 and 15 (Total 90 min)<br><u>Active lever presses</u><br>Two-way ANCOVA<br>Between-subjects factors: Abstinence day, M+B dose<br>Covariate: Inactive (90 min) | Abstinence day (1,15) between-subjects<br>M+B dose (0, 50+50μg/side) between-subjects<br>Abstinence day X M+B dose interaction | $F_{1,43}$ = 43.4<br>$F_{1,43}$ = 10.5<br>$F_{1,43}$ = 7.3 | <0.001*<br>0.002*<br>0.01* | 0.502<br>0.197<br>0.144 |
| Figure 2D. Relapse tests days 1 and 15 (90 min time course)<br><u>Active lever presses</u><br>Mixed ANCOVA<br>Between-subjects factors: Abstinence day, M+B dose<br>Within-subjects factor: Session time<br>Covariate: Inactive (90 min) | Abstinence day (1,15) between-subjects<br>M+B dose (0, 50+50μg/side) between-subjects<br>Abstinence day X M+B dose interaction<br><br>Session time (30, 60, 90) within-subjects<br>Session time X Abstinence day interaction<br>Session time X M+B dose interaction<br>Session time X Abstinence day X M+B dose interaction | $F_{1,43}$ = 43.3<br>$F_{1,43}$ = 10.5<br>$F_{1,43}$ = 7.3<br><br>$F_{2,86}$ = 62.7<br>$F_{2,86}$ = 14.4<br>$F_{2,86}$ = 4.4<br>$F_{2,86}$ = 2.7 | <0.001*<br>0.002*<br>0.01*<br><br><0.001*<br><0.001*<br>0.015*<br>0.073 | 0.502<br>0.197<br>0.145<br><br>0.593<br>0.251<br>0.093<br>0.059 |

**Exp. 3.** Effect of muscimol-baclofen vSub inactivation on incubation after forced abstinence

| Figure number | Factor name | F-value | *p*-value | Partial Eta$^2$ |
|---|---|---|---|---|
| Figure 3B. Self-administration training<br>Infusions<br>RM-ANOVA<br>Within-subjects factor: Session | Session (1-14) within-subjects | $F_{13,429}$ = 20.0 | <0.001* | 0.377 |
| Figure 3B. Self-administration training<br>Active Lever presses<br>RM-ANOVA<br>Within-subjects factors: Session, Lever | Session (1-14) within-subjects<br>Lever (Active, Inactive) within subjects<br>Lever X Session interaction | $F_{13,429}$ = 4.8<br>$F_{1,33}$ = 48.6<br>$F_{13,429}$ = 5.4 | <0.001*<br><0.001*<br><0.001* | 0.128<br>0.596<br>0.140 |
| Figure 3C. Relapse test day 15 (Total 90 min)<br>Active lever presses<br>One-Way ANCOVA<br>Between-subjects factor: M+B dose<br>Covariate: Inactive (90 min) | M+B dose (0, 50+50µg/side) between-subjects | $F_{1,31}$ = 0.2 | 0.644 | 0.007 |
| Figure 3C. Relapse test day 15 (90 min time course)<br>Active lever presses<br>Mixed ANCOVA<br>Between-subjects factor: M+B dose<br>Within-subjects factor: Session time<br>Covariate: Inactive (90 min) | M+B dose (0, 50+50µg/side) between-subjects<br>Session time (30, 60, 90) within-subjects<br>Session time X M+B dose interaction | $F_{1,31}$ = 0.2<br>$F_{2,62}$ = 30.2<br>$F_{2,62}$ = 0.4 | 0.644<br><0.001*<br>0.671 | 0.007<br>0.494<br>0.013 |

**Exp. 4.** Effect of Daun02 vSub inactivation on incubation after electric barrier-induced abstinence

| Figure number | Factor name | F-value | *p*-value | Partial Eta$^2$ |
|---|---|---|---|---|
| Figure 4B. Self-administration training<br>Infusions<br>RM-ANOVA<br>Within-subjects factor: Session | Session (1-14) within-subjects | $F_{13,754}$ = 85.2 | <0.001* | 0.595 |
| Figure 4B. Self-administration training<br>Active lever presses<br>RM-ANOVA<br>Within-subjects factors: Session, Lever | Session (1-14) within-subjects<br>Lever (Active, Inactive) within subjects<br>Lever X Session interaction | $F_{13,754}$ = 14.8<br>$F_{1,58}$ = 55.8<br>$F_{13,754}$ = 14.6 | <0.001*<br><0.001*<br><0.001* | 0.203<br>0.490<br>0.201 |
| Figure 4C. Electric barrier<br>Infusions<br>RM-ANOVA<br>Within-subjects factor: Session | Session (1-13) within-subjects | $F_{12,696}$ = 139.1 | <0.001* | 0.706 |
| Figure 4C. Electric barrier<br>Active lever presses<br>RM-ANOVA<br>Within-subjects factors: Session, Lever | Session (1-13) within-subjects<br>Lever (Active, Inactive) within subjects<br>Lever X Session interaction | $F_{12,696}$ = 27.6<br>$F_{1,58}$ = 11.4<br>$F_{12,696}$ = 33.5 | <0.001*<br>0.001*<br><0.001* | 0.323<br>0.164<br>0.366 |
| Figure S1. SA context Induction session day 15 (15 min) | Daun02 dose (0, 4µg/side) between-subjects | $F_{1,23}$ = 0.02 | 0.896 | 0.001 |

| | | | | |
|---|---|---|---|---|
| Figure 4D. SA context Relapse test day 18 (Total 90 min)<br><u>Active lever presses</u><br>One-Way ANCOVA<br>Between-subjects factor: Daun02 dose<br>Covariate: Inactive (90 min) | Daun02 dose (0, 4µg/side) between-subjects | $F_{1,23} = 4.5$ | 0.044* | 0.165 |
| Figure 4D. SA context Relapse tests day 18 (90 min time course)<br><u>Active lever presses</u><br>Mixed ANCOVA<br>Between-subjects factor: Daun02 dose<br>Within-subjects factor: Session time<br>Covariate: Inactive (90 min) | Daun02 dose (0, 4µg/side) between-subjects<br>Session time within-subjects<br>Session time X Daun02 dose interaction | $F_{1,23} = 4.5$<br>$F_{2,46} = 20.9$<br>$F_{2,46} = 1.1$ | 0.044*<br><0.001*<br>0.332 | 0.165<br>0.476<br>0.047 |
| Figure 4E. SA context vSub Fos-Xgal immunohistochemistry<br><u>Fos and Xgal counts</u><br>One-way ANOVA<br>Between-subjects factor: Daun02 dose | Fos: Daun02 dose (0, 4µg/side) between-subjects<br>Xgal: Daun02 dose (0, 4µg/side) between-subjects | $F_{1,24} = 50.3$<br>$F_{1,24} = 33.1$ | <0.001*<br><0.001* | 0.677<br>0.580 |
| Figure 4D. Novel context Relapse test day 18 (Total 90 min)<br><u>Active lever presses</u><br>One-Way ANCOVA<br>Between-subjects factor: Daun02 dose<br>Covariate: Inactive (90 min) | Daun02 dose (0, 4µg/side) between-subjects | $F_{1,30} = 0.08$ | 0.779 | 0.003 |
| Figure 4D. Novel context Relapse tests day 18 (90 min time course)<br><u>Active lever presses</u><br>Mixed ANCOVA<br>Between-subjects factor: Daun02 dose<br>Within-subjects factor: Session time<br>Covariate: Inactive (90 min) | Daun02 dose (0, 4µg/side) between-subjects<br>Session time within-subjects<br>Session time X Daun02 dose interaction | $F_{1,30} = 0.08$<br>$F_{2,60} = 39.1$<br>$F_{2,60} = 0.8$ | 0.779<br><0.001*<br>0.46 | 0.003<br>0.566<br>0.026 |
| Figure 4E. Novel context vSub Fos-Xgal immunohistochemistry<br><u>Fos and Xgal counts</u><br>One-way ANOVA<br>Between-subjects factor: Daun02 dose | Fos: Daun02 dose (0, 4µg/side) between-subjects<br>Xgal: Daun02 dose (0, 4µg/side) between-subjects | $F_{1,31} = 0.5$<br>$F_{1,31} = 0.3$ | 0.491<br>0.618 | 0.015<br>0.008 |

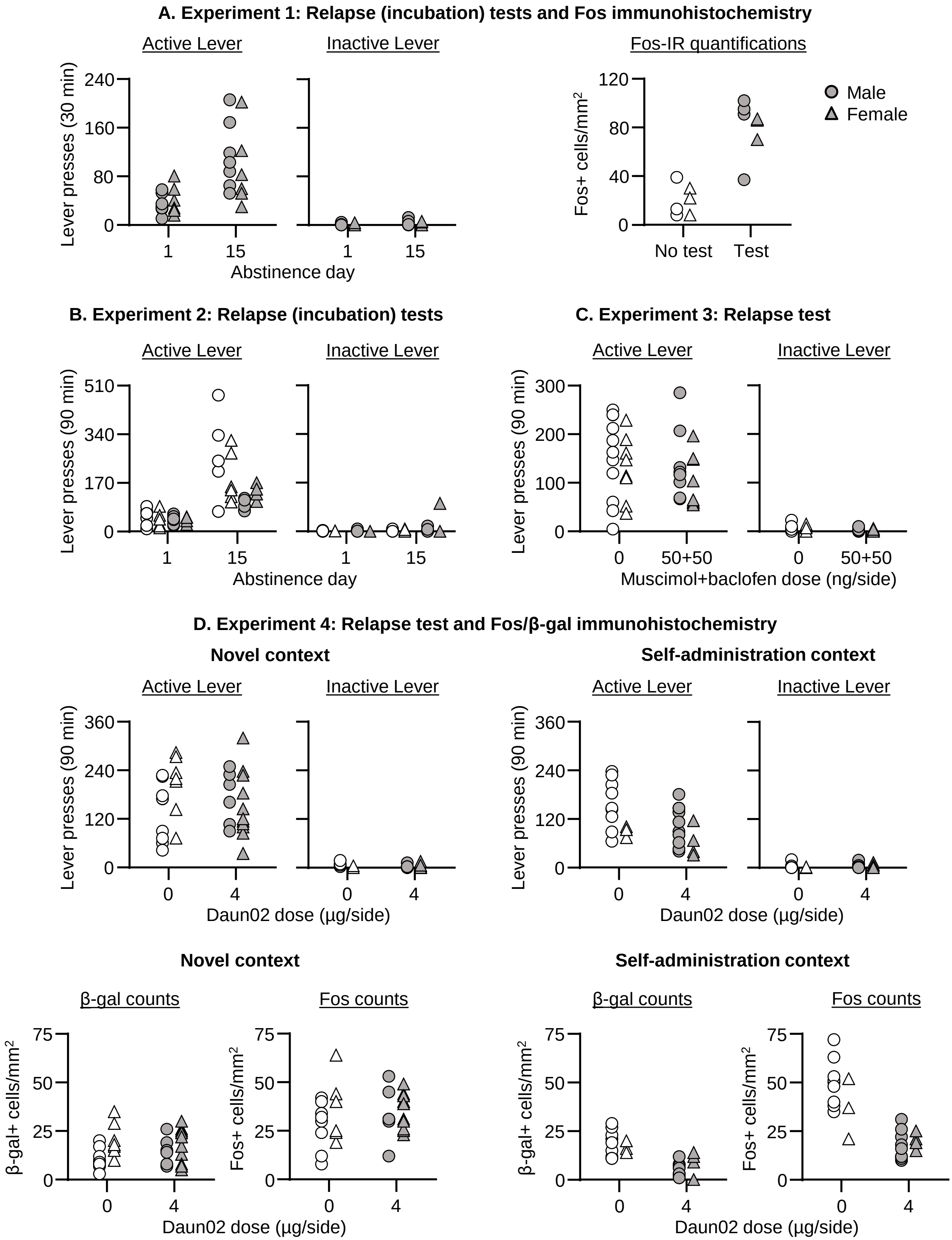


**Figure S1.** **(A-C)** Number of active and inactive lever presses during the relapse tests of individual rats in Exp. 1-3, and Fos+ cells (counts/mm$^2$) of individual rats in Exp. 1. **(D)** Number of active and inactive lever presses during the relapse tests and number of β-gal+ or Fos+ cells (counts/mm$^2$) of individual rats in Exp. 4.
